# Gene model for the ortholog of *Ilp3* in *Drosophila pseudoobscura*

**DOI:** 10.64898/2026.08.19.745830

**Authors:** Bethany C. Lieser, Leon F. Laskowski, Robyn Huber, Katheryn Opperman Kolker, Andrew M Arsham, Chinmay P Rele, Stephanie Toering Peters

## Abstract

Gene model for the ortholog of *Insulin-like peptide 3* (*Ilp3*) in the D. pseudoobscura Apr. 2013 (BCM-HGSC Dpse_3.0/DpseGB3) Genome Assembly (GenBank Accession: GCA_000001765.2) of *Drosophila pseudoobscura*. This ortholog was characterized as part of a developing dataset to study the evolution of the Insulin/insulin-like growth factor signaling pathway (IIS) across the genus *Drosophila* using the Genomics Education Partnership gene annotation protocol for Course-based Undergraduate Research Experiences.

## Introduction

*This article reports a predicted gene model generated by undergraduate work using a structured gene model annotation protocol defined by the Genomics Education Partnership (GEP; thegep.org) for Course-based Undergraduate Research Experience (CURE). The following information may be repeated in other articles submitted by participants using the same GEP CURE protocol for annotating Drosophila species orthologs of Drosophila melanogaster genes in the insulin signaling pathway*.

“In this GEP CURE protocol students use web-based tools to manually annotate genes in non-model *Drosophila* species based on orthology to genes in the well-annotated model organism fruitfly *Drosophila melanogaster*. The GEP uses web-based tools to allow undergraduates to participate in course-based research by generating manual annotations of genes in non-model species (Rele et al., 2023). Computational-based gene predictions in any organism are often improved by careful manual annotation and curation, allowing for more accurate analyses of gene and genome evolution (Mudge and Harrow 2016; Tello-Ruiz et al., 2019). These models of orthologous genes across species, such as the one presented here, then provide a reliable basis for further evolutionary genomic analyses when made available to the scientific community.” (Myers et al., 2024).

“The particular gene ortholog described here was characterized as part of a developing dataset to study the evolution of the Insulin/insulin-like growth factor signaling pathway (IIS) across the genus *Drosophila*. The Insulin/insulin-like growth factor signaling pathway (IIS) is a highly conserved signaling pathway in animals and is central to mediating organismal responses to nutrients (Hietakangas and Cohen 2009; Grewal 2009).” (Myers et al., 2024).

“Invertebrate insulins function similarly to metazoan insulin-like growth factors and play a role in cell and organ growth (Chan 2000). In *Drosophila*, seven insulin-like peptides (Ilp1-Ilp7) have a two-chain structure similar to vertebrate insulin and interact with the sole insulin-like receptor, InR, to initiate the insulin signaling cascade (Brogiolo et al., 2001; Nassel and Vanden Broeck 2016). Like the *Ilp2* and *Ilp5* genes, the *Ilp3* gene is expressed in median neurosecretory cells (MNCs) in the brain (Ikeya et al., 2002). While the seven Ilps act redundantly with respect to promoting growth, they also have unique expression patterns and functions (Ikeya et al., 2002; Grönke et al., 2010). Ilp3 may act with the transcription factor dFOXO in a positive feedback loop to regulate Ilp2 and Ilp5 secretion from MNCs (Grönke et al., 2010). In female *Drosophila*, ablation of MNCs or knockout of *Ilp3* have been shown to reduce fecundity and remating rates (Grönke et al., 2010; Wigby et al., 2011). Knockout of *Ilp3* also results in sleep defects (Yamaguchi et al., 2022).” (Gruys et al., 2024).

“*D. pseudoobscura* is part of the *pseudoobscura* species subgroup within the *obscura* species group in the subgenus *Sophophora* of the genus *Drosophila* (Sturtevant 1942; Buzzati-Traverso and Scossiroli 1955). It was first described by Frolova (1929). The *pseudoobscura* species subgroup is endemic to the western hemisphere, where *D. pseudoobscura* is distributed throughout Western North America, Mexico, and Central America (Markow and O’Grady 2006). An additional population of *D. pseudoobscura*, found near Bogota, Colombia, is partially reproductively isolated from the North and Central American populations (Prakash 1972). *D. pseudoobscura* is found primarily in chaparral and temperate forests. *D. pseudoobscura* has been studied extensively in the context of ecological and behavioral genetics, speciation, and genome evolution (Powell 1997).” (Lawson et al., 2025).

We propose a gene model for the *D. pseudoobscura* ortholog of the *D. melanogaster Insulin-like peptide 3* (*Ilp3*) gene. The genomic region of the ortholog corresponds to the uncharacterized protein XP_002135097.1 (Locus ID LOC6900737) in the *D. pseudoobscura* Apr. 2013 (BCM-HGSC Dpse_3.0/DpseGB3) Genome Assembly of *D. pseudoobscura* (GCA_000001765.2; *Drosophila* 12 Genomes Consortium, 2007; Richards et al., 2005). This model is based on RNA-Seq data from

*D. pseudoobscura* (SRP006203; Graveley et al., 2011) and *Ilp3* in *D. melanogaster* using FlyBase release FB2024_02 (GCA_000001215.4; Gramates et al., 2022; Jenkins et al., 2022; Larkin et al., 2021).

## Results

### Synteny

The target gene, *Ilp3*, occurs on chromosome 3L in *D. melanogaster* and is nested within *CG32052* alongside *Insulin-like peptide 4* (*Ilp4*) and *Insulin-like peptide 2* (*Ilp2*). *Ilp3* is flanked upstream by *Inhibitor-2* (*I-2*) and *CG43897*, which nests *Insulin-like peptide 5* (*Ilp5*), and downstream by *Insulin-like peptide 1* (*Ilp1*) and *Z band alternatively spliced PDZ-motif protein 67* (*Zasp67*). The *tblastn* search of *D. melanogaster* Ilp3-PA (query) against the *D. pseudoobscura* (GenBank Accession: GCA_000001765.2) Genome Assembly (database) placed the putative ortholog of *Ilp3* within scaffold CH379069.3 at locus LOC6900737 (XP_002135097.1) with an E-value of 3e-10 and a percent identity of 39.71%. Furthermore, the putative ortholog is nested by LOC4812906 (XP_015043245.1) alongside LOC4813890 (XP_001353874.3) and LOC117185171 (XP_033241287.1), which correspond to *CG32052, Ilp4*, and *Ilp2* in *D. melanogaster* (E-value: 0.0, 2e-28, and 1e-34; identity: 84.45%, 51.20%, and 46.58%, respectively, as determined by *blastp*; Figure 1A, Altschul et al., 1990). The putative ortholog is flanked upstream by LOC4813105 (XP_001353871.2) and LOC6900735 (XP_033241266.1), which nests LOC6900736 (XP_002135096.3), and correspond to *I-2, CG43897*, and *Ilp5* in *D. melanogaster* (E-value: 8e-81, 0.0, and 8e-17; identity: 79.81%, 62.47%, and 35.71%, respectively, as determined by *blastp*). The putative ortholog of *Ilp3* is flanked downstream by LOC4813355 (XP_001353876.4) and LOC4813356 (XP_033241280.1), which correspond to *Ilp1* and *Zasp67* in *D. melanogaster* (E-value: 2e-38 and 0.0; identity: 57.02% and 71.41%, respectively, as determined by *blastp*). The putative ortholog assignment for *Ilp3* in *D. pseudoobscura* is supported by the following evidence: The genes surrounding the *Ilp3* ortholog are orthologous to the genes at the same locus in *D. melanogaster* and synteny is completely conserved, supported by e-values and percent identities, so we conclude that LOC6900737 is the correct ortholog of *Ilp3* in *D. pseudoobscura* (Figure 1A).

**Figure 1:**
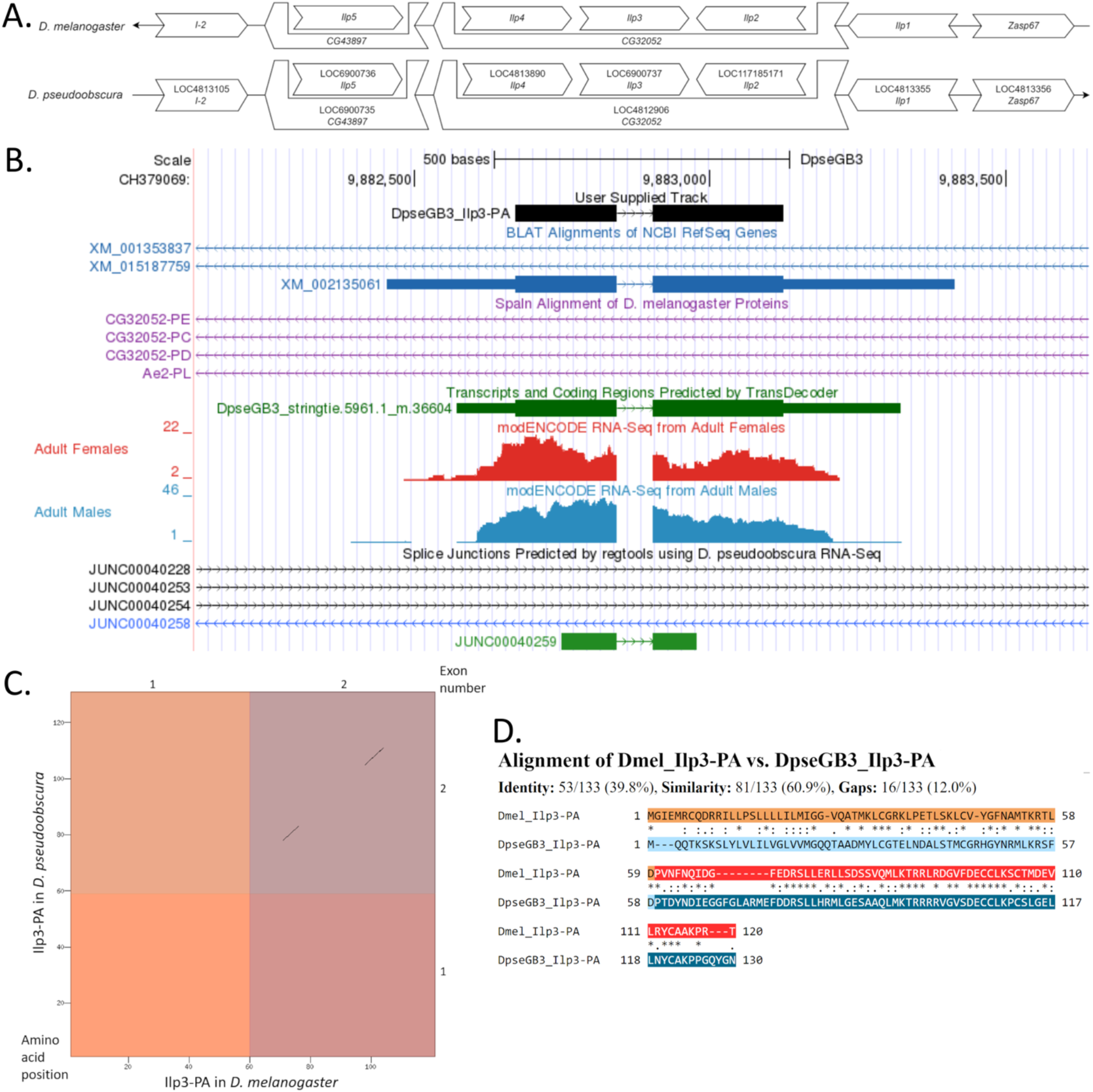
*Ilp3* gene model comparison between *Drosophila pseudoobscura* and *Drosophila melanogaster*. **(A) Synteny comparison of the genomic neighborhoods for *Ilp3* in *Drosophila melanogaster* and *D. pseudoobscura***. Thin underlying arrows indicate the DNA strand within which the target gene–*Ilp3*–is located in *D. melanogaster* (top) and *D. pseudoobscura* (bottom). The thin arrow pointing to the right indicates that *Ilp3* is on the positive (+) strand in *D. pseudoobscura*, and the thin arrow pointing to the left indicates that *Ilp3* is on the negative (-) strand in *D. melanogaster*. The wide gene arrows pointing in the same direction as *Ilp3* are on the same strand relative to the thin underlying arrows, while wide gene arrows pointing in the opposite direction of *Ilp3* are on the opposite strand relative to the thin underlying arrows. White gene arrows in *D. pseudoobscura* indicate orthology to the corresponding gene in *D. melanogaster*. Gene symbols given in the *D. pseudoobscura* gene arrows indicate the orthologous gene in *D. melanogaster*, while the locus identifiers are specific to *D. pseudoobscura*. **(B) Gene Model in GEP UCSC Track Data Hub** (Raney et al., 2014). The coding-regions of *Ilp3* in *D. pseudoobscura* are displayed in the User Supplied Track (black); CDSs are depicted by thick rectangles and introns by thin lines with arrows indicating the direction of transcription. Subsequent evidence tracks include BLAT Alignments of NCBI RefSeq Genes (dark blue, alignment of Ref-Seq genes for *D. pseudoobscura*), Spaln of D. melanogaster Proteins (purple, alignment of Ref-Seq proteins from *D. melanogaster*), Transcripts and Coding Regions Predicted by TransDecoder (dark green), RNA-Seq from Adult Females and Adult Males (red and light blue respectively; alignment of Illumina RNA-Seq reads from *D. pseudoobscura*; Chen et al., 2014), and Splice Junctions Predicted by regtools using *D. pseudoobscura* RNA-Seq (SRP006203, Graveley et al., 2011). Splice junction (JUNC00040259), shown in green, has a read-depth of 50. **(C) Dot Plot of Ilp3-PA in *D. melanogaster* (*x*-axis) vs. the orthologous peptide in *D. pseudoobscura* (*y*-axis)**. Amino acid number is indicated along the left and bottom; CDS number is indicated along the top and right, and CDSs are also highlighted with alternating colors. **(D) Protein alignment between *D. melanogaster* Ilp3-PA and its putative ortholog in *D. pseudoobscura***. The alternating colored rectangles represent adjacent CDSs. The symbols in the match line denote the level of similarity between the aligned residues. An asterisk (*) indicates that the aligned residues are identical. A colon (:) indicates the aligned residues have highly similar chemical properties—roughly equivalent to scoring > 0.5 in the Gonnet PAM 250 matrix (Gonnet et al., 1992). A period (.) indicates that the aligned residues have weakly similar chemical properties— roughly equivalent to scoring > 0 and £ 0.5 in the Gonnet PAM 250 matrix. A space indicates a gap or mismatch when the aligned residues have no similarity—roughly equivalent to scoring £ 0 in the Gonnet PAM 250 matrix.

### Protein Model

*Ilp3* in *D. pseudoobscura* has two CDSs within the genome sequence. The protein sequence (Ilp3-PA) is translated from one mRNA isoform (*Ilp3-RA*; Figure 1B). Relative to the ortholog in *D. melanogaster*, the CDS number and protein isoform-count are conserved. The sequence of Ilp3-PA in *D. pseudoobscura* has 49.49% identity (E-value: 7e-28) with the protein-coding isoform Ilp3-PA in *D. melanogaster*, as determined by *blastp* (Figure 1C). Coordinates of this curated gene model are stored by NCBI at GenBank/BankIt (accession **BK065280**).

## Methods

“Detailed methods including algorithms, database versions, and citations for the complete annotation process can be found in Rele et al. (2023). Briefly, students use the GEP instance of the UCSC Genome Browser v.435 (https://gander.wustl.edu; Kent WJ et al., 2002; Navarro Gonzalez et al., 2021) to examine the genomic neighborhood of their reference IIS gene in the *D. melanogaster* genome assembly (Aug. 2014; BDGP Release 6 + ISO1 MT/dm6). Students then retrieve the protein sequence for the *D. melanogaster* target gene for a given isoform and run it using *tblastn* against their target *Drosophila* species genome assembly on the NCBI BLAST server (https://blast.ncbi.nlm.nih.gov/Blast.cgi, Altschul et al., 1990) to identify potential orthologs. To validate the potential ortholog, students compare the local genomic neighborhood of their potential ortholog with the genomic neighborhood of their reference gene in *D. melanogaster*. This local synteny analysis includes at minimum the two upstream and downstream genes relative to their putative ortholog. They also explore other sets of genomic evidence using multiple alignment tracks in the Genome Browser, including BLAT alignments of RefSeq Genes, Spaln alignment of D. melanogaster proteins, multiple gene prediction tracks (e.g., GeMoMa, Geneid, Augustus), and modENCODE RNA-Seq from the target species.

Genomic structure information (e.g., CDSs, intron-exon number and boundaries, number of isoforms) for the *D. melanogaster* reference gene is retrieved through the Gene Record Finder (https://gander.wustl.edu/~wilson/dmelgenerecord/index.html; Rele et al., 2023). Approximate splice sites within the target gene are determined using *tblastn* using the CDSs from the *D. melanogaste*r reference gene. Coordinates of CDSs are then refined by examining aligned modENCODE RNA-Seq data, and by applying paradigms of molecular biology such as identifying canonical splice site sequences and ensuring the maintenance of an open reading frame across hypothesized splice sites. Students then confirm the biological validity of their target gene model using the Gene Model Checker (https://gander.wustl.edu/~wilson/dmelgenerecord/index.html; Rele et al., 2023), which compares the structure and translated sequence from their hypothesized target gene model against the *D. melanogaster* reference gene model. At least two independent models for this gene were generated by students under mentorship of their faculty course instructors. These models were then reconciled by a third independent researcher mentored by the project leaders to produce the final model presented here. Note: comparison of 5’ and 3’ UTR sequence information is not included in this GEP CURE protocol.” (Gruys et al., 2025)

## Acknowledgements

We would like to thank Wilson Leung for developing and maintaining the technological infrastructure that was used to create this gene model and Laura K. Reed for overseeing the project. Thank you to FlyBase for providing the definitive database for *Drosophila melanogaster* gene models. Further, we would like to thank the editors and developers at the journal *microPublication: Biology* for assistance in developing the template for these single gene ortholog publications.

## Funding

This material is based upon work supported by the National Science Foundation (1915544) and the National Institute of General Medical Sciences of the National Institutes of Health (R25GM130517) to the Genomics Education Partnership (GEP; https://thegep.org/; PI-LKR). Any opinions, findings, and conclusions or recommendations expressed in this material are solely those of the author(s) and do not necessarily reflect the official views of the National Science Foundation nor the National Institutes of Health.

## Notes

### Competing Interest Statement

The authors have declared no competing interest.

